# Patterns of retrieval-related cortico-striatal connectivity are stable across the adult lifespan

**DOI:** 10.1101/2022.03.16.484376

**Authors:** Paul F. Hill, Marianne de Chastelaine, Michael D. Rugg

## Abstract

Memory retrieval effects in the striatum are well documented and robust across a number of experimental paradigms and stimulus materials. However, the functional significance of these effects, and whether they are moderated by age, remains unclear. In the present study, we used fMRI paired with an associative recognition task to examine retrieval effects in the striatum in a large sample of healthy young, middle aged, and older adults. We identified anatomically segregated patterns of enhanced striatal BOLD activity during recollection- and familiarity-based memory judgments. Successful recollection was associated with enhanced BOLD activity in bilateral putamen and nucleus accumbens. Familiarity effects were evident in the head of the caudate nucleus bilaterally. Crucially, none of these effects were reliably moderated by age. Using psychophysiological interaction analyses, we observed a monitoring-related increase in functional connectivity between the caudate and regions of the frontoparietal control network, and between the putamen and bilateral retrosplenial cortex and intraparietal sulcus. In all instances, monitoring-related increases in cortico-striatal connectivity were unmoderated by age. These results suggest that the striatum, and the caudate in particular, couples with the frontoparietal control network to support top-down retrieval monitoring operations, and that the integrity of these effects are preserved in advanced age.

## 1. Introduction

Memory retrieval effects in the striatum are well documented and robust across a number of experimental paradigms and stimulus materials (for reviews, see Kim, 2011; Scimeca & Badre, 2012; Spaniol et al., 2009). The striatum underlies a number of goal-directed behaviors, including motor planning, cognitive control, and complex value judgments. In real-world environments, accurate memory retrieval confers a number of economic, social, and survival advantages and can therefore be conceived of as a goal-directed behavior. However, the functional significance of striatal retrieval effects remain ambiguous and a matter of ongoing debate. Retrieval effects have been reported in several anatomically distinct subregions of the striatum ranging across its ventral and dorsal aspects (Clos et al., 2015; Elward et al., 2015; King et al., 2018). In the present study, we examined functionally differentiated patterns of cortico-striatal connectivity during successful memory retrieval as a means to disentangle the functional significance of striatal retrieval effects.

Early models of retrieval success effects in the striatum focused on the motivational significance of making a correct decision in a memory test. Support for such models came from an early study investigating the impact of monetary incentives associated either with accurate recognition of previously studied items (associative hits) or with detection of unstudied new items (correct rejections) (Han et al., 2010; Herweg et al., 2018). Enhanced activity in bilateral caudate was evident for both types of memory judgments when they were associated with reward. In light of these findings, Han and colleagues (2010) proposed that retrieval success effects in the striatum reflect perceived goal attainment, rather than successful memory retrieval. Consequently, striatal retrieval effects in the absence of experimental reward manipulations were held to reflect the intrinsic value associated with detecting previously studied items (Han et al., 2010).

Contrary to the unitary ‘goal-attainment’ account described by Han and colleagues (2010), a previous study demonstrated dissociable reward and episodic retrieval effects in the striatum (Elward et al., 2015). Participants performed a source memory task in which an accurate source memory judgment was associated with either high or low monetary reward (depending on which of the two possible contexts was retrieved), while inaccurate judgments were associated with a commensurate monetary loss. Accurate source judgments, but not reward expectation, were associated with enhanced activity in the left lateral putamen whereas reward, but not retrieval, effects were evident in the ventral striatum. These results converge with findings from several other studies to indicate that retrieval and reward effects in the striatum reflect functionally distinct processes and thus cannot be accounted for by a unitary construct such as goal attainment (see also Clos et al., 2015; Schwarze et al., 2013).

Alternative accounts of retrieval-related striatal activity suggest that, particularly when coupled with the prefrontal cortex, the striatum supports top-down cognitive control of memory retrieval (Buckner, 2004; Scimeca & Badre, 2012). Following a retrieval attempt, controlled monitoring processes are engaged to evaluate the contents of retrieval in relation to behavioral goals and strategies (Burgess & Shallice, 1996; Rugg & Wilding, 2000). The striatum has been suggested to support controlled memory retrieval by updating and prioritizing retrieved content so as to increase the likelihood of retrieving goal-relevant details in support of a memory judgment, while simultaneously damping the influence of goal-irrelevant details (i.e., adaptive gating) (Scimeca & Badre, 2012). These types of controlled memory processes are generally assumed to be vulnerable to advancing age (Braver & Barch, 2002; Buckner, 2004; Grady, 2012; Hedden & Gabrieli, 2004), which may in turn hinder the ability to align recollected content with behavioral goals (e.g., Srokova et al., 2021). One possibility, which we will address in this study, is that age-related decline in the functional integrity of cortico-striatal networks result in reduced recollection accuracy.

Finally, and of particular relevance to the present study, King and colleagues (2018) reported evidence of anatomically dissociable striatal effects during recollection- and familiarity-driven memory judgments. Enhanced BOLD activity in bilateral caudate nucleus was evident for both recollected and familiar test items. A second bilateral cluster in the nucleus accumbens, extending into subgenual frontal cortex, was exclusively sensitive to recollection. Of importance, the results reported by King and colleagues (2018) converged across three independent studies that employed different operationalizations of recollection and familiarity (source memory, associative recognition, remember/know tests) and both verbal and non-verbal materials, suggesting that recollection- and familiarity-driven effects in the striatum are invariant with respect to these factors.

Patterns of cortico-striatal connectivity estimated with resting-state fMRI provide additional context for exploring retrieval effects in the striatum (Buckner, 2004; Choi et al., 2012; Fjell et al., 2016; Gordon et al., 2021; Nyberg et al., 2016; Rieckmann et al., 2018). Endogenous functional connectivity between the dorsal caudate and regions comprising the frontoparietal control network has been reported to correlate with performance on offline tests of episodic memory (Buckner, 2004; Rieckmann et al., 2018). Similar associations between memory performance and functional coupling between ventral aspects of the striatum (including ventral caudate and nucleus accumbens) and regions of the default mode network ((Di Martino et al., 2008)) and hippocampus (Nyberg et al., 2016; Postuma & Dagher, 2006) have also been observed. Along with the ventral striatum, the hippocampus and regions of the default mode network are consistently engaged during successful episodic retrieval, underscoring the possibility that the ventral striatum is a component of the so-called ‘core recollection network’ (Rugg & Vilberg, 2013). Whether a similar functional topography extends to task-evoked changes in cortico-striatal connectivity during an online memory retrieval task is unclear.

In the present study, we examined striatal retrieval effects in a large sample of healthy young, middle aged, and older adults. Participants studied visually presented word pairs followed by an associative recognition memory test during fMRI scanning. Test items on the associative recognition test comprised pairs of ‘intact’ (studied together), ‘rearranged’ (studied on different trials), and ‘new’ (unstudied) word pairs. We addressed three principal questions: 1) Are retrieval effects evident in anatomically dissociable striatal subregions in a large, lifespan sample of adults? 2) Are any such retrieval effects in the striatum moderated by age? And 3) do retrieval success effects in the striatum elicit dissociable patterns of recollection- and familiarity-driven functional connectivity with other cortical and subcortical regions? To address the latter question, we employed psychophysiological interaction (PPI) analyses to identify regions of enhanced functional connectivity with striatal subregions during recollection and familiarity judgments.

## 2. Methods & Materials

Data from the encoding (de Chastelaine et al., 2016a) and retrieval (de Chastelaine et al., 2016b, 2017) phases of the present study have been described in previous publications. The principal findings reported below comparing retrieval-related striatal functional connectivity across age groups have not previously been reported. More detailed descriptions of participant demographics, experimental procedures, and behavioral results are reported elsewhere (de Chastelaine et al., 2016a; 2016b; 2017).

### 2.1. Participants

Participants were 36 younger adults (18-29 yrs.; *M* = 22 yrs; *SD* = 3.0 yrs; 17 females), 36 middle aged adults (43–55 yrs.; *M*=49 yrs.; *SD*=3.4 yrs.; 17 female), and 64 older adults (63-76 yrs.; *M* = 68 yrs.; *SD* = 3.6 yrs.; 35 females). Participant were recruited from the University of Texas at Dallas and surrounding communities. All participants gave informed consent in accordance with the UT Dallas and University of Texas Southwestern Institutional Review Boards and were compensated at the rate of $30 per hour. All participants were right-handed and were fluent in English before the age of five. No participant had a history of neurological or psychiatric disease, substance abuse, diabetes, or untreated hypertension. None were taking prescription medication that affected the central nervous system. All participants completed a neuropsychological test battery on a day prior to the experimental MRI session and these results are reported elsewhere (de Chastelaine et al., 2016b).

### 2.2. Materials

Experimental stimuli comprised 320 semantically unrelated word pairs randomly separated into four lists of 80 pairs. For each set of yoked participants (one young, one middle-aged, and one or two older participants), word pairs from three of the lists were pseudo-randomly ordered to form the study list. The test list included 320 critical word pairs divided into 160 ‘intact’ pairs (words presented together at study), 80 ‘rearranged’ pairs (words presented on different trials at study, and 80 ‘new’ pairs (words not encountered at study. An additional 60 word pairs were used as practice stimuli. Critical test pairs were intermixed with 106 null trials (white fixation). Two filler pairs were placed at the start and two in the middle of each study and test block (see below), and a 30 s rest break occurred midway through each block. In both the study and test phases, word pairs were presented for 2 s with inter-item intervals of 1.5 s (study) and 2.5 s (test). Cogent 2000 software (www.vislab.ucl.ac.uk/cogent.php) implemented in Matlab 2012b (www.mathworks.com) was used for stimulus presentation and response collection.

### 2.3. Experimental Procedure

Participants were given instructions and practice sessions for both the study task and the memory test prior to scanning. The study and test phases took place in separate scanning sessions. During study, participants indicated with a button press which of the two objects denoted by the words in each pair was more likely to fit into the other. Study pairs were presented in two consecutive blocks separated by an approximately 2 min inter-block interval. After completing the study task, participants excited the scanner for a brief break.

After a 15 min break participants reentered the scanner to complete an associative memory test that was presented in three consecutive blocks separated by 2 min inter-block intervals. The memory test required one of three keyed responses to indicate whether each test pair was intact, rearranged, or new. An ‘intact’ response was required when participants recognized both words and had a specific memory of the two words having been presented together at study. Participants were instructed not to guess and to only respond ‘intact’ only when they were reasonably confident that the words had been studied together. A ‘rearranged’ response was required when both words were recognized from the study phase but there was no specific memory of the words having been paired together previously. Participants were instructed to make a ‘new’ response when neither word, or only one word, was recognized.

### 2.4. MRI Acquisition and Analysis

Functional and anatomical images were acquired with a Philips Achieva 3T MR scanner (Philips Medical System, Andover, MA USA) equipped with a 32 channel parallel imaging head coil. A T1-weighted anatomical image was acquired with a 3D MPRAGE pulse sequence (FOV-256×224, voxel size 1×1×1 mm, 160 slices, sagittal acquisition). Functional scans were acquired with a T2*-weighted EPI with the following parameters: TR 2 s, TE 30 ms, flip angle 70 deg, field-of-view 240×240, matrix size 80×78). Each EPI volume comprised 33 slices (3mm thickness, 1mm inter-slice gap) with an in-plane resolution of 3×3mm. Slices were acquired in ascending order, oriented parallel to the AC-PC line and positioned for full coverage of the cerebrum and most of the cerebellum. The functional data were acquired using a sensitivity encoding (SENSE) reduction factor of 2. fMRI data were acquired during both study and test phases. The first five volumes of each block were discarded to allow tissue magnetization to achieve a steady state. Test sessions were concatenated to form a single time-series prior to model estimation.

Functional and anatomical images were preprocessed using SPM8 (Wellcome Department of Cognitive Neurology, London, UK), run under Matlab R2008a (MathWorks). Univariate and PPI analyses were carried out using SPM12 run under Matlab R2017a. Functional images were motion and slice-time corrected, realigned, and spatially normalized using a sample-specific template (see de Chastelaine et al., 2015, for further details) based on the MNI reference brain. Images were resampled into 3mm isotropic voxels and smoothed with an isotropic 8mm full-width half-maximum Gaussian kernel. T1-weighted anatomical images were normalized with a procedure analogous to that applied to the functional images. For each participant, item-elicited neural activity was modeled using a delta function convolved with the canonical hemodynamic response function (HRF).

#### 2.4.1. GLM Analysis

The fMRI data from the associative memory test were analyzed in two stages. At the first stage, a separate GLM was constructed for each participant. Three events of interest were included in the design matrix: correctly endorsed intact pairs (*associative hits*), intact pairs incorrectly identified as rearranged (*associative misses*), and correctly endorsed new pairs (*correct rejections*). Each event of interest was modeled with a delta function convolved with SPM’s canonical hemodynamic response function (HRF). Intact and rearranged pairs wrongly judged as new, correctly endorsed rearranged pairs, false alarms, filler trials, and trials where no response was given were each modeled as covariates of no interest, along with six regressors representing motion-related variance (three for rigid-body translation and three for rotation), and constants representing means across each scan session. Data from volumes showing a transient displacement of >1mm or >1° in any direction were eliminated by defining them as covariates of no interest. The time series in each voxel were high-pass filtered to 1/128 Hz to remove low-frequency noise and scaled within session to a grand mean of 100 across voxels and scans.

Parameters estimates from the three events of interest (associative hits, associative misses, correct rejections) were carried forward to a 3 (age group) x 3 (item type) mixed-design ANOVA as implemented within SPM12 (and hence employing a single pooled error term). Recollection effects were operationalized as greater BOLD activity for associative hits than for associative misses. The reverse contrast was employed to operationalize retrieval monitoring (associative misses > associative hits). Familiarity effects were operationalized as greater BOLD activity for associative misses (i.e. recognized but un-recollected word pairs) than for correctly endorsed unstudied pairs (correct rejections). The reverse contrast (correct rejections > associative misses) was employed to operationalize novelty effects. See de Chastelaine et al. 2016a and 2017 for theoretical justification of these contrasts.

The principal GLM analyses were conducted using contrasts derived from the ANOVA model (height-threshold *p* < .001, uncorrected). Family-wise error (FWE) corrected cluster-extent thresholds were estimated with the Gaussian random field method implemented in SPM12. To test *a priori* predictions regarding striatal recognition effects, we applied small volume corrections for each contrast within an anatomically defined bilateral basal ganglia mask. The mask was created by combining the bilateral accumbens, caudate, and putamen labels from the Neuromorphometrics atlas provided in SPM12. For each significant striatal cluster (p < .05 FWE corrected), we extracted parameter estimates for the BOLD responses elicited by the respective test responses (associative hits, associative misses, correct rejections), averaged across all voxels falling within a 3 mm radius of the peak voxel. Results of whole brain analyses for this study have been previously described in detail elsewhere (de Chastelaine et al., 2016; 2017).

#### 2.4.2. Psychophysiological Interaction Analyses

We performed psychophysiological interaction (PPI) analyses to identify memory-related modulation of functional connectivity. PPI analyses were conducted using six striatal seed regions-of-interest (ROIs) which were identified by the recollection and familiarity contrasts (see below ‘3.1. Univariate GLM Results’): left and right caudate nucleus, left and right putamen, and left and right nucleus accumbens (NAc). The loci of the six striatal seed regions were derived separately for each participant. Using the mass univariate effects reported above, the peak recollection (left and right NAc, left and right putamen) and familiarity (left and right caudate) effect falling within a 10 mm radius of the striatal coordinates listed in Table 1 were identified for each participant. The seeds were then defined as all voxels falling within a 3 mm radius sphere centered on each peak.

**Table 1.** Loci of peak striatal effects.

| Region | Contrast | MNI |  |  | Peak-z | Cluster Size |
| --- | --- | --- | --- | --- | --- | --- |
|  |  | x | y | z |  |  |
| L. PUT | Recollection | -18 | -4 | 22 | 4.34 | 21 |
| R. PUT | Recollection | 18 | -7 | 25 | 6.16 | 38 |
| L. NAc | Recollection | -12 | 8 | -14 | Inf | 159 |
| R. NAc | Recollection | 12 | 8 | -11 | Inf | 75 |
| L. CN | Familiarity | -12 | 11 | 1 | Inf | 48 |
| R. CN | Familiarity | 15 | 14 | 4 | Inf | 53 |

For each seed ROI and principal memory contrast (recollection, familiarity), regression models were created at the individual subject level that included three regressors of interest: a physiological regressor, a psychological (task) regressor, and a PPI regressor. The physiological variable was the representative time course (first eigenvariate) of all voxels falling within each seed ROI. The psychological regressors were constructed separately for recollection (associative hit > associative miss) and familiarity (associative miss > correct rejection) contrasts. For the recollection condition, psychological regressors were constructed by creating a vector that coded associative hit trials as 1, associative misses as -1, and all other trials as 0. The psychological regressors for the familiarity condition were constructed similarly, with associative misses coded as 1, correct rejections coded as -1, and all other trials as 0. The respective task vectors were then convolved with the canonical HRF in SPM12 to create the psychological regressor included in the first-level models. To create the PPI regressor of interest, the physiological regressor was deconvolved from the HRF, multiplied by the unconvolved task regressor, and then reconvolved with the HRF.

To identify memory-related changes in functional connectivity, we ran separate first-level PPI analyses for each of the seed regions (left and right caudate, left and right NAc, left and right putamen) and for two contrasts of interest (associative hits > associative misses; associative misses > correct rejections). As described above, the reversal of these contrasts corresponds to retrieval monitoring and novelty effects, respectively. Patterns of recollection- and familiarity-driven connectivity change (positive PPI effects) could thus be dissociated from patterns of monitoring- and novelty-driven connectivity change (negative PPI effects).

First-level PPI contrast images were carried forward to separate second-level 3 (age) x 2 (hemisphere) factorial ANOVAs for the respective contrasts (recollection, familiarity) and striatal subregions (caudate, putamen, NAc). Whole brain analyses were conducted using contrasts derived from the ANOVA models. For each group-level model, we used two-sided F-contrasts (height threshold *p* < .001 uncorrected, clusterwise FWE *p* < .05) to identify the main effects of age, seed region, hemisphere (left, right) as well as the age x hemisphere interaction. Family-wise error (FWE) corrected cluster-extent thresholds were estimated with the Gaussian random field method implemented in SPM12.

## 3. Results

### 3.1. Univariate GLM results

Regions of enhanced striatal BOLD activity associated with the recollection and familiarity contrasts are illustrated in Figure 1 and reported in Table 1. Recollection effects were evident in bilateral putamen and NAc. For both subregions, recollection effects did not differ significantly by hemisphere or age group. Average parameter estimates collapsed across hemisphere and age group did not differ between associative misses and correct rejections but were enhanced for recollected test items (Figure 1B, middle and right panels). The familiarity contrast revealed enhanced BOLD activity in the head of the caudate bilaterally. Familiarity effects in the caudate also did not differ by hemisphere or age group. Parameter estimates collapsed across hemisphere and age group did not differ between associative hits and associative misses but were significantly attenuated for correct rejections (Figure 1B, left panel). The retrieval monitoring and novelty contrasts did not reveal any enhanced striatal BOLD effects.

**Figure 1.**
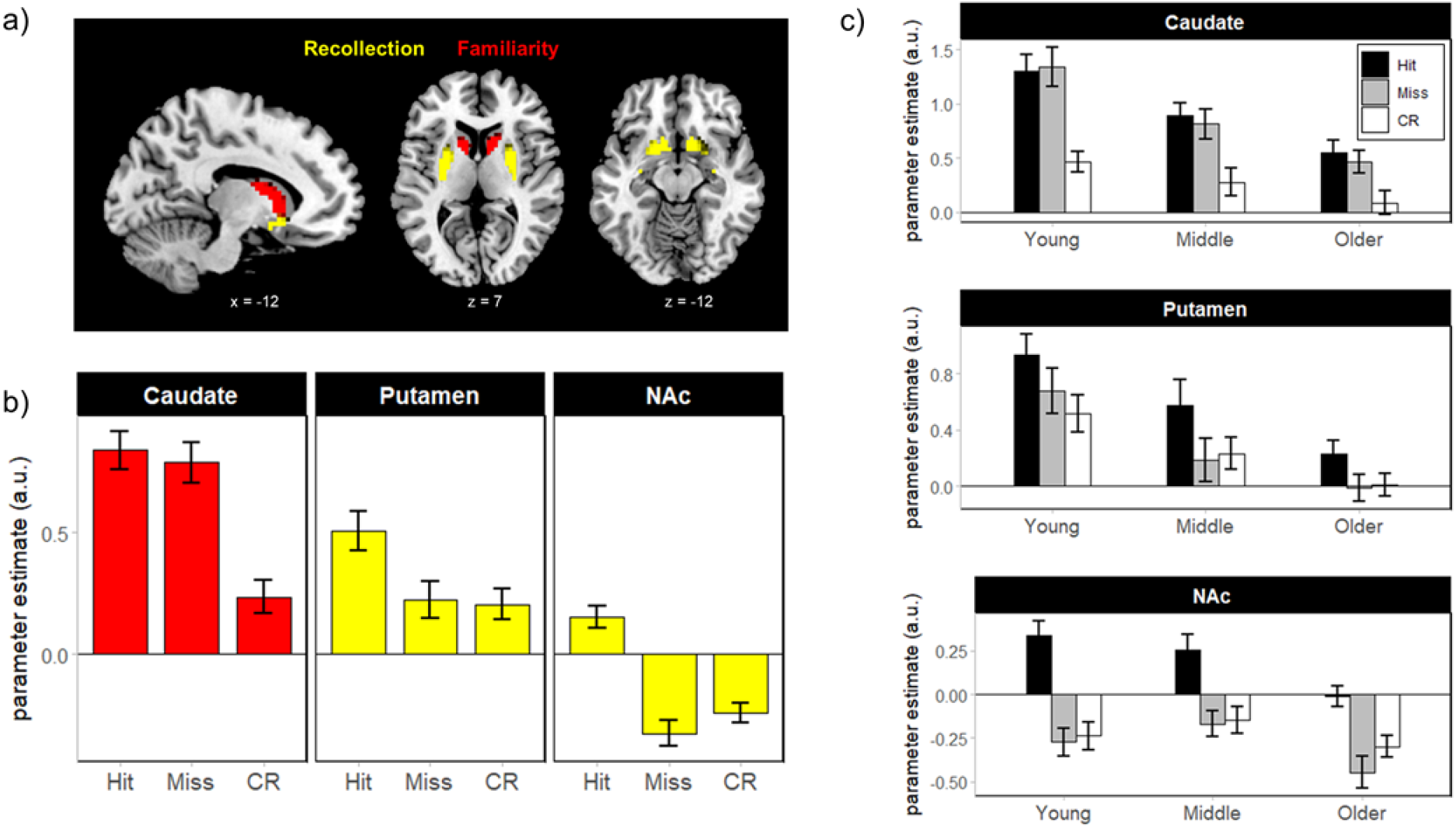
Mean recollection and familiarity striatal BOLD effects. a) Clusters demonstrating across-age recollection (yellow) and familiarity (red) effects. b) Mean parameter estimates for associative hits, associative misses, and correct rejections (CR), collapsed across age group and hemisphere. c) Mean parameter estimates plotted for each age group. NAc = nucleus accumbens. Error bars reflect standard error of the mean.

Visual inspection of Figure 1C suggests a robust main effect of age group, such that striatal responses decline with age. To confirm this impression, we submitted the parameter estimates extracted from each striatal subregion (collapsed across hemisphere) to separate 3 (age group: young, middle-aged, older) x 3 (memory: associative hit, associative miss, correct rejection) ANOVAs. These analyses confirmed that none of the recollection and familiarity effects differed with age (age x memory interaction *ps* > .07). However, significant main effects of age were evident in each of the striatal subregions [caudate: *F*(1,133) = 10.77, *p* < .001, partial-η^2^ = .139; putamen: *F*(1,133) = 7.94, *p* < .001, partial-η^2^ = .107; NAc: *F*(1,133) = 3.86, *p* = .023, partial-η^2^ = .055]. Follow-up pairwise comparisons revealed that caudate activity (collapsed across memory conditions) was greater in the young adult group relative to the older adult group (*t*(133) = -4.62, *p* < .001). There were no significant differences between caudate activity in the middle-aged group and either of the other groups (*ps* > .06). An identical pattern was evident in the putamen, with greater activity in the young adults relative older adults (*t*(133) = -3.98, *p* < .001) and no significant differences between putamen activity in the middle-aged group and either of the other groups (*ps* > .09). Finally, activity in the NAc was greater in middle-aged adults compared to older adults (*t*(133) = 2.48, *p* = .038). Activity in the NAc did not significantly differ between younger adults and either of the other two age groups (*ps* > .1).

### 3.2. Memory-related changes in striatal connectivity

To determine whether the magnitude of memory-related changes in functional connectivity with striatal subregions varied across age groups, we entered first-level parameter estimates of the PPI regressor into a group-level 3 (age: young, middle, older) x 2 (hemisphere) factorial ANOVA separately for each contrast (recollection, familiarity) and striatal subregion (caudate, putamen, NAc). We failed to identify any brain regions that demonstrated a significant main effect of age group or hemisphere on the magnitude of memory-related changes in connectivity (two-sided F-contrast), even at relaxed statistical thresholds (*p* < .01, uncorrected). As a result, for each of the four second-level analyses we report the results of one-sided *t*-contrasts collapsed across age group and hemisphere.

#### 3.2.1. Monitoring-related connectivity change

Contrary to our *a priori* predictions, we did not identify any brain regions where successful recollection (associative hits > associative misses) was associated with increased connectivity with any of the striatal subregions. Instead, the retrieval monitoring contrast (i.e., the inverse of the recollection contrast) revealed several clusters that exhibited enhanced connectivity with the caudate for associative misses compared to associative hits, including anterior cingulate, left ventrolateral prefrontal cortex, right dorsolateral prefrontal cortex, and left intraparietal sulcus (Figure 2A). The retrieval-monitoring contrast also revealed enhanced connectivity between the putamen and a large cluster with a peak voxel in the left caudate and extending into a number of posterior brain regions. Increasing the height threshold (*p* < .0005) revealed separate clusters in the left caudate body, left and right retrosplenial cortex, and left and right intraparietal sulcus (Figure 2B). We did not identify any regions of enhanced recollection- or monitoring-related NAc functional connectivity. A complete list of regions showing enhanced monitoring-related striatal connectivity is provided in Table 2.

**Table 2.**
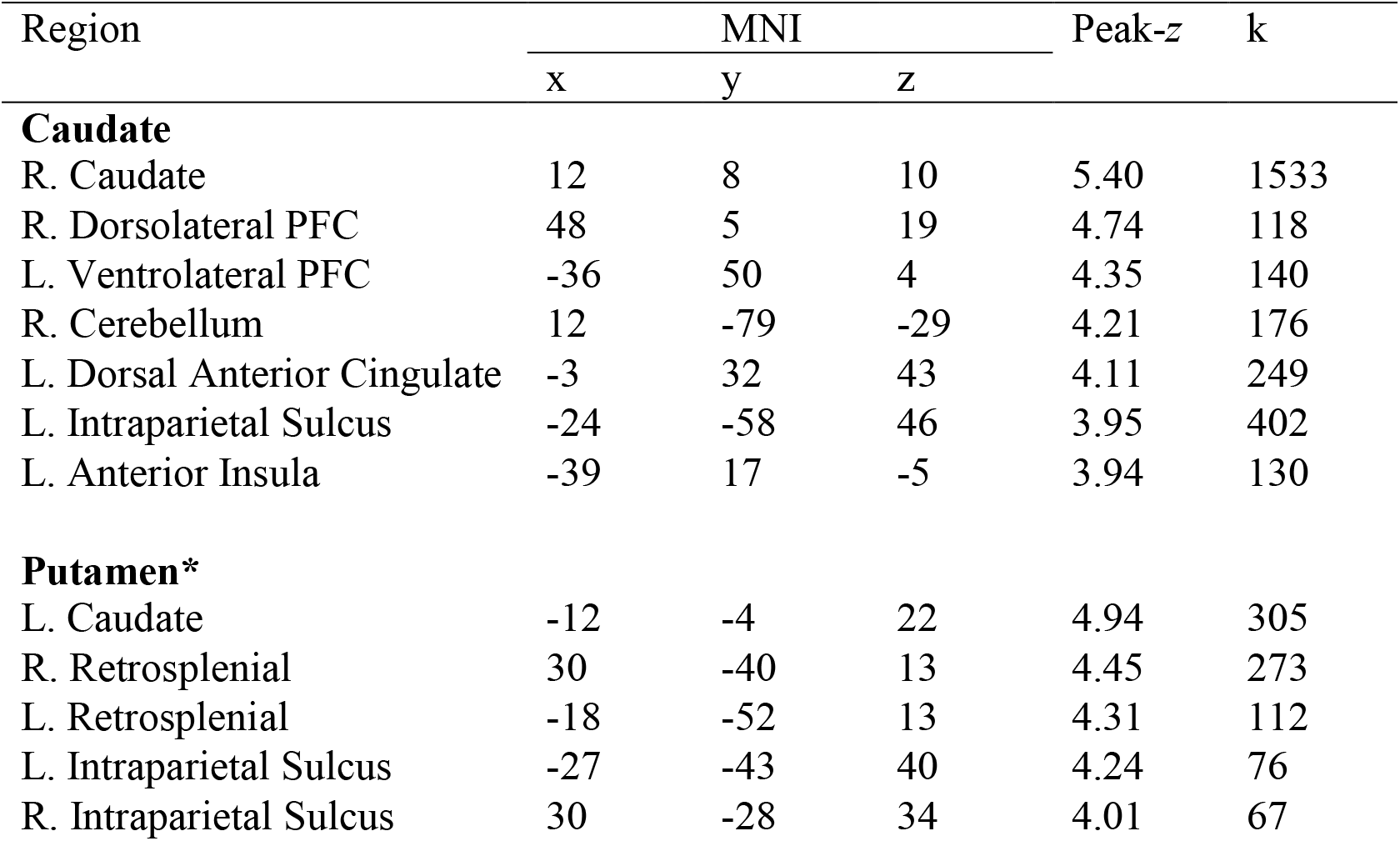
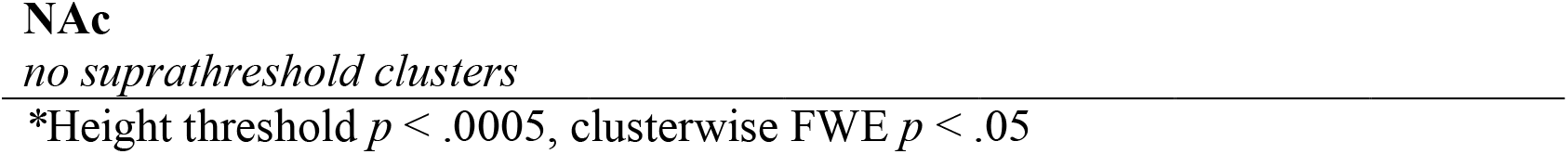
Peak loci of monitoring-driven connectivity change.

| Region | MNI |  |  | Peak-z | k |
| --- | --- | --- | --- | --- | --- |
|  | x | y | z |  |  |
| **Caudate** |  |  |  |  |  |
| R. Caudate | 12 | 8 | 10 | 5.40 | 1533 |
| R. Dorsolateral PFC | 48 | 5 | 19 | 4.74 | 118 |
| L. Ventrolateral PFC | -36 | 50 | 4 | 4.35 | 140 |
| R. Cerebellum | 12 | -79 | -29 | 4.21 | 176 |
| L. Dorsal Anterior Cingulate | -3 | 32 | 43 | 4.11 | 249 |
| L. Intraparietal Sulcus | -24 | -58 | 46 | 3.95 | 402 |
| L. Anterior Insula | -39 | 17 | -5 | 3.94 | 130 |
| **Putamen\*** |  |  |  |  |  |
| L. Caudate | -12 | -4 | 22 | 4.94 | 305 |
| R. Retrosplenial | 30 | -40 | 13 | 4.45 | 273 |
| L. Retrosplenial | -18 | -52 | 13 | 4.31 | 112 |
| L. Intraparietal Sulcus | -27 | -43 | 40 | 4.24 | 76 |
| R. Intraparietal Sulcus | 30 | -28 | 34 | 4.01 | 67 |

**Figure 2.**
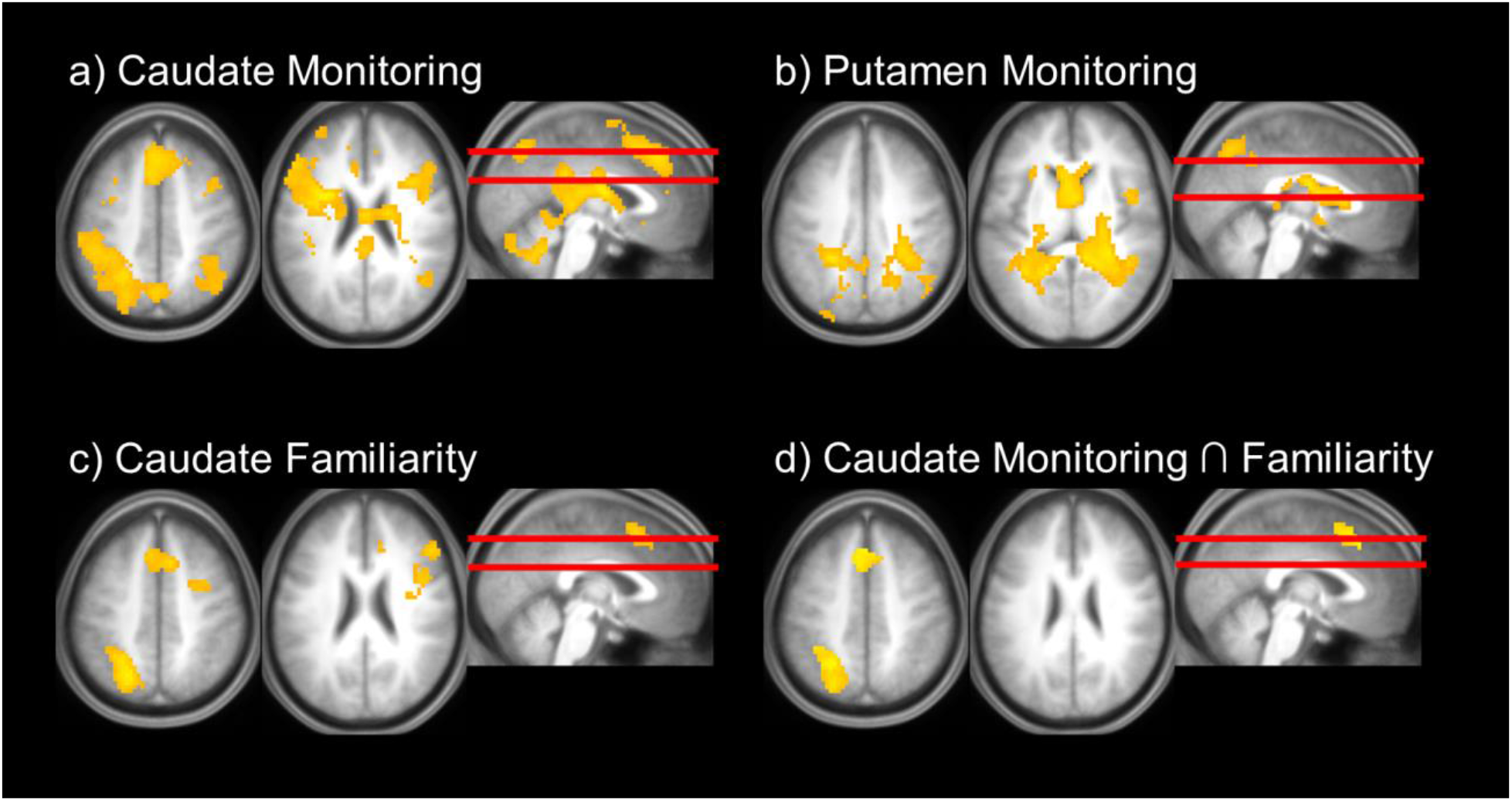
Striatal psychophysiological interaction (PPI) results. Enhanced monitoring-related connectivity with (a) bilateral head of the caudate nucleus and (b) bilateral putamen. (c) Enhanced familiarity-related connectivity with head of the caudate nucleus. (d) Overlap between enhanced monitoring- and familiarity-related caudate connectivity. Images displayed at height threshold *p* < .005.

#### 3.2.2. Familiarity-related connectivity change

Turning to the familiarity contrast (associative misses > correct rejections), we identified a pattern of increased connectivity between the caudate and clusters in the dorsomedial prefrontal cortex (extending into dorsal anterior cingulate), left intraparietal sulcus, and bilateral occipital cortex (Table 3). An additional cluster in the right dorsolateral prefrontal cortex was also evident at a less conservative height threshold (*p* < .005, clusterwise FWE *p* < .05). As can be seen in Figure 2A and 2C, this pattern of familiarity-driven connectivity appeared to overlap with regions demonstrating enhanced caudate monitoring-related connectivity. To confirm this impression, we inclusively masked the main effect of monitoring-driven caudate connectivity with the corresponding main effect of familiarity-driven caudate connectivity (mask threshold *p* < .005, uncorrected). As can be seen in Figure 2D, this procedure confirmed that patterns of enhanced familiarity- and monitoring-driven connectivity change with the caudate overlapped in regions comprising the frontoparietal control network, including anterior cingulate (xyz = -3, 32, 43, *z* = 4.11, *k* = 116) and left intraparietal sulcus (xyz = -24, -58, 46, *z* = 3.95, *k* = 145). We did not identify any familiarity-related changes in connectivity with the NAc or putamen. Nor did we observe any regions of enhanced novelty-related connectivity change with any of the striatal subregions.

**Table 3.** Peak loci of familiarity-driven connectivity change.

| Region | MNI |  |  | Peak-z | k |
| --- | --- | --- | --- | --- | --- |
|  | x | y | z |  |  |
| **Caudate** |  |  |  |  |  |
| L. Intraparietal Sulcus | -21 | -67 | 43 | 4.64 | 175 |
| R. Pre-SMA/ACC | 12 | 17 | 52 | 4.41 | 113 |
| L. Lateral Occipital | -33 | -100 | -2 | 4.24 | 184 |
| R. Lateral Occipital | 30 | -94 | -2 | 4.04 | 102 |
| R. Dorsolateral PFC* | 48 | 8 | 28 | 3.85 | 292 |

### 3.3. Overlap between striatal connectivity and mean BOLD activity

We next examined the extent to which patterns of monitoring- and familiarity-related striatal connectivity change overlapped with the mass univariate BOLD effects identified by the recollection (associative hits > associative misses), retrieval monitoring (associative misses > associative hits), familiarity (associative misses > correct rejections), and novelty (correct rejections > associative misses) contrasts. We accomplished this by inclusively masking the respective group-level PPI *t*-contrasts (uncorrected height threshold *p* < .001, clusterwise FWE *p* < .05) with each of the mean univariate retrieval contrasts (mask threshold *p* < .001) in turn.

Regions of enhanced monitoring-related connectivity change with the caudate overlapped with mean BOLD monitoring effects in anterior cingulate cortex (xyz = -3, 32, 43; *z* = 4.11; *k* = 180) and right dorsolateral prefrontal cortex (xyz = 48, 8, 19; *z* = 4.15; *k* = 264; PPI height threshold *p* < .005, clusterwise FWE *p* < .05). A similar pattern was observed for the familiarity contrasts; enhanced familiarity-driven connectivity changes overlapped with mean familiarity effects in the anterior cingulate (xyz = 12, 17, 52; *z* = 4.41; *k* = 111) and left intraparietal sulcus (xyz = -21, -67, 43; *z* = 4.64; *k* = 153). We did not observe any overlap between enhanced caudate connectivity and BOLD recollection or novelty effects. Nor did patterns of enhanced monitoring-related putamen connectivity overlap with any of the mean monitoring effects.

## 4. Discussion

We used fMRI to examine memory retrieval effects in the striatum in a large sample of healthy young, middle aged, and older adults. Consistent with prior findings, we identified anatomically segregated patterns of enhanced striatal BOLD activity during recollection- and familiarity-based recognition judgments. In the head of the caudate, both recollected and familiar test items elicited BOLD responses that were indistinguishable from one another and were greater than the responses elicited by novel test pairs. By contrast, recollected test items elicited BOLD responses in the NAc and putamen that were reliably greater than responses elicited by familiar and novel test items. Our findings are highly consistent with previous reports of dissociations between recollection- and familiarity-related fMRI BOLD responses in ventral and dorsal aspects of the striatum in young adults (King et al., 2018; note that the study reported by King et al. included the young adult participants from this study and therefore does not represent a fully independent analysis). The inter-regional dissociations observed in the present study add to existing evidence that retrieval effects in the striatum reflect multiple, functionally distinct cognitive processes (Clos et al., 2015; Elward et al., 2015; King et al., 2018; Schwarze et al., 2013). Of importance, the present fMRI BOLD retrieval effects in the striatum were seemingly impervious to age. These null findings are consistent with prior reports of absent or very limited effects of age in studies of episodic memory retrieval (e.g. Wang et al, 2016; Hou et al., 2021; de Chastelaine et al 2016b; 2017). There were, however, robust main effects of age in each striatal region. This dissociation between retrieval effects and main effects is strongly reminiscent of that observed in retrieval-sensitive cortical regions, where null effects of age on recollection and familiarity effects have also been reported to co-exist with significant age differences in mean BOLD activity (e.g. Hou et al., 2022, Wang et al., 2016).

Using PPI, we sought to identify regions demonstrating retrieval-related modulation of functional connectivity with the striatum. Within the caudate, we observed overlapping patterns of connectivity with cortical regions often reported in association with familiarity-based memory judgments and top-down post-retrieval monitoring (cf. Buckner, 2004; Gordon et al., 2021; Rieckmann et al., 2018; see Rugg, in press for review). Of importance, we note that the respective contrasts that gave rise to these effects, familiarity and retrieval monitoring, share associative misses as a common condition. Associative misses are held to reflect the conjunction of an above criterion item-driven familiarity signal and a failure to recollect associative information (de Chastelaine et al. 2016). They are therefore assumed to impose heavier demands on post-retrieval monitoring than correctly endorsed intact pairs (associative hits) or new pairs (correct rejections) by virtue of the need to resolve a conflict between the familiarity of the individual test items and a weak or absent recollection signal (de Chastelaine et al., 2016b; Horne et al., 2021; see also Achim & Lepage, 2005). Thus, whereas the caudate responds generally to the perceived ‘oldness’ of a test item, the presence of an ambiguous recollection signal appears to enhance frontal-striatal coupling. We suggest that increased functional connectivity between the caudate and regions comprising the frontoparietal control network reflects the heightened demands placed by items attracting associative misses on executive control processes supporting evaluation of the outcome of a retrieval attempt in relation to behavioral goals.

In contrast to the higher order cognitive and motivational functions of the caudate and NAc, respectively, cortico-striatal circuits involving the putamen have traditionally been associated with motor functions (Alexander, 1986; Gordon et al., 2021). However, our observation of robust recollection effects in the putamen converges with findings from studies reporting retrieval success effects in this region that are difficult to explain by a motoric account (Elward et al., 2015; Koster et al., 2015). Here, we observed enhanced functional connectivity between the putamen and retrosplenial cortex, albeit in response to associative misses relative to successfully recollected test items. On first sight, this finding seems at odds with the presence of reliable recollection effects in both the putamen and bilateral retrosplenial cortex. One possibility is that test trials endorsed as familiar elicited a negative striatal prediction error reflecting the absence of an above-criterion recollection signal. When retrieved content deviates from expectations, error signals can adjust retrieval strategies to increase the likelihood of retrieving relevant information (Scimeca & Badre, 2012). Negative recollection signals in the putamen may mobilize engagement of on-line memory search operations and, consequently, modulate activity in regions of the core recollection network such as retrosplenial cortex. Although speculative, this interpretation is consistent with recent evidence that striatal reward prediction errors modulate episodic and semantic memory (Calderon et al., 2021; Ergo et al., 2020; Pine et al., 2018; Scimeca et al., 2016; Sinclair & Barense, 2019).

Age did not significantly moderate the strength of monitoring-related connectivity between the caudate and frontoparietal control network or between the putamen and bilateral clusters in retrosplenial cortex and intraparietal sulcus. This age-invariant pattern of task-evoked functional coupling contrasts with findings from a recent study that reported reduced resting-state functional connectivity between the caudate and regions of the frontoparietal control network in older adults, accompanied by less *decoupling* between the caudate and regions of the default mode network (Rieckmann et al., 2018; see also Fjell et al., 2016; Nyberg et al., 2016). The present findings might suggest that task-evoked and resting-state measures of striatal connectivity capture separate (albeit spatially overlapping) inter-regional processes that are differentially impacted by age (Geerligs et al., 2015; Laumann & Snyder, 2021; Parker & Razlighi, 2019). This interpretation is of course speculative and additional research comparing resting-state and task-evoked cortico-striatal connectivity will be necessary to adjudicate between this and other possibilities.

One limitation of the present study is our use of a cross-sectional approach to examine age differences in retrieval-related cortico-striatal connectivity. This type of cross-sectional approach fails to account for the potential impact of longitudinal aging processes on age-related variance in both memory performance and the neural correlates of recognition memory judgments (Nyberg et al., 2010; Rugg, 2016). Nevertheless, the present results demonstrate that patterns of retrieval-related cortico-striatal connectivity are stable across healthy young, middle-aged, and older adult cohorts.

That being said, our conclusion that age does not impact retrieval-related changes in cortico-striatal connectivity rests on the assumption that the transfer function mediating the relationship between neural activity and the fMRI BOLD signal is age- and regionally-invariant. There is however evidence that cerebro-vascular reactivity (CRV), a driver of the BOLD signal, declines with advancing age (e.g., Lu et al., 2011). Future studies correcting for age differences in CVR may therefore yield different results.

In conclusion, we identified anatomically segregated patterns of enhanced striatal BOLD activity during recollection- and familiarity-based memory judgments that were seemingly not moderated by age. Using PPI, we observed a pattern of increased functional connectivity between the head of the caudate and regions of the frontoparietal control network often associated with post retrieval monitoring. Recollection success effects were evident in bilateral putamen and retrosplenial cortex, though functional connectivity between these regions was maximal for familiar test items eliciting a sub-criterion recollection signal. Importantly, retrieval success effects in the striatum, as well as all patterns of task-evoked changes in functional connectivity, were also unmoderated by age, suggesting that the cortico-striatal circuits supporting memory retrieval remain stable across much of the adult lifespan.

## Acknowledgements

This project was supported by the National Institute on Aging (grant number 1RF1AG039103).

## Notes

**Conflict of Interest:** None

### Competing Interest Statement

The authors have declared no competing interest.

